# Detectability of biotin tags by LC-MS/MS

**DOI:** 10.1101/2020.12.30.424786

**Authors:** Lorenz Nierves, Philipp F. Lange

## Abstract

The high affinity of biotin to streptavidin has made it one of the most widely used affinity tags in proteomics. Early methods used biotin for enrichment alone and mostly ignored the biotin labeled peptide. Recent advances in labeling led to an increase in biotinylation efficiency and shifted the interest to detection of the site of biotinylation. This increased confidence in identification and provides additional structural information yet it requires efficient release of the biotinylated protein/peptide and sensitive separation and detection of biotinylated peptides by LC-MS/MS. Despite its long use in affinity proteomics the effect of biotinylation on the chromatographic, ionization, and fragmentation behaviour and ultimate detection of peptides is not well understood. To address this we compare two commercially-available biotin labels EZ-Link Sulfo-NHS-Biotin and Sulfo-NHS-SS-Biotin, the latter one containing a labile linker to efficiently release biotin to determine the effects of peptide modification on peptide detection. We describe an increase of hydrophobicity and charge reduction with increasing number of biotin labels attached. Based on our data we recommend gradient optimization to account for more hydrophobic biotinylated peptides and include singly charged precursors to account for charge reduction by biotin.

## INTRODUCTION

Biotin is an often-used affinity tag in many proteomic workflows. Its stable non-covalent interaction with its known binding partner, (strept)avidin^1–3^, has been exploited to interrogate protein localization (Cell Surface Capturing technology^4^), synthesis (PUNCH-P^5^) and protein interaction (BioID^6^). Traditionally most biotin-based labeling and enrichment approaches relied on indirect identification enrichment of biotinylated proteins and detection of enriched tryptic peptides. Direct detection of the biotinylated peptide has only recently become more popular for protein interaction (BioSite^7^, DiDBiT^8^) and post-translational modification (subtiligase-based neo-N termini^9^) mapping as it provides direct evidence of the target protein and, more importantly, the site of biotinylation. With increased efficiency of the biotinylation reactions the number of biotin labels added to the protein of interest and their impact on the physicochemical properties of proteins and peptides increases. To date the potential effects of different biotin tags on mass spectrometric detection are only incompletely understood. Here we study two frequently used commercial biotin tags to guide optimal liquid chromatography and mass spectrometer configuration and enable accurate and complete identification of biotinylated peptides.

To enable detection of biotinylated peptides following enrichment, they first need to be released from their affinity matrix. While release from anti-biotin antibodies at low pH is straightforward, efficient reversal of the high-affinity (K_d_=∼10^−15^) interaction with streptavidin poses a challenge. One attractive solution is the introduction of a cleavable disulfide bridge linker between target protein and biotin moiety. This allows for release of the biotinylated protein or peptide in reducing conditions.

Typical acquisition by mass spectrometer relies on positive charges from the N terminus and C terminal lysine or arginine residue that resulted from trypsin digestion. However, because of its reactive amine group the same lysine is often the target for chemical modification including biotinylation. Modification of the lysine side chain can alter its ability to retain a positive charge and thus alter ionization, fragmentation and detection. Similarly, liquid chromatography is commonly optimized for primarily hydrophilic tryptic peptides and chemical adducts may alter hydrophobicity and chromatographic behavior of peptides. Here we hypothesize that the biotin tag significantly alters the physicochemical properties and detectability of peptides and that different biotin tags can fundamentally differ in their behaviour and suitability for MS detection.

We compare two commercially-available biotin labels: EZ-Link Sulfo-NHS-Biotin (biotin-NHS; Thermo Scientific, cat. no.: 21217) and EZ-Link Sulfo-NHS-SS-Biotin (biotin-SS-NHS; Thermo Scientific, cat. no.: 21331) to determine the effects of peptide modification on peptide detection. Both reagents are water-soluble and possess the NHS ester moiety to facilitate biotinylation of primary amines. Biotin-NHS has a short linker, whereas biotin-SS-NHS has a longer linker that includes a disulfide bridge to release the captured protein from (strept)avidin in reducing conditions. We further characterize the effects of biotin modification compared to unlabeled peptides and suggest changes to chromatography as mass spectrometry methods to account for biotinylation induced changes.

## MATERIALS AND METHODS

### Biotin labeling of HeLa peptides

10-15 µg prepared HeLa peptides were mixed with 100mM HEPES, pH=8.5 at a 1:2 (v/v) ratio. The pH of the resulting solution was confirmed to be around pH=8.0-8.5. The commercially-available biotin was prepared in a stock solution of 20mM using 100mM HEPES, pH 8.5 as the diluant. The biotin was added to the HeLa peptides to a final concentration of 2mM biotin. For unlabeled peptides, an equivalent volume of 100mM HEPES, pH 8.5 was added. Samples were incubated on ice for 30 minutes. Labeling reaction was quenched using 50mM Tris-HCl, pH 8.0. For the reduction and alkylation of the biotin-NHS-SS label, a final concentration of 10mM DTT and 50mM CAA was added, respectively. During reduction, samples were incubated at 37°C for 30 minutes; alkylation was conducted at room temperature (in the dark) for 30 minutes.

### Preparation of STaGE tips

Samples were prepared for LC-MS/MS analysis or other downstream assays using STaGE tips as prepared in Rappsilber et al. (2003)^12^. Briefly, two small circular EmporeTM SPE C18 disks were punched out using a flat-end needle. A straightened paper clip was used to position the disks into a P200 pipette tips. STaGE tips were conditioned with 40 µL methanol, 40 µL 0.1% Formic Acid (FA), 60% Acetonitrile (ACN), and 40 µL 0.1% Trifluoroacetic Acid (TFA). Samples pH was adjusted to pH=2.0–3.0 using 10% TFA prior to loading. Samples were eluted with 40 µL 0.1% FA, 60% ACN. The ACN was eliminated from the samples via SpeedVac. Samples were re-suspended in 0.1% FA.

### LC-MS/MS analysis

Peptide concentration and total were determined using the Nanodrop. A total of 1 µg peptide per sample was injected for analysis. Mass spectrometric analyses were performed on Q Exactive HF Orbitrap mass spectrometer coupled with an Easy-nLC 1200 liquid chromatography system (Themo Scientific). Buffer A was 2% Acetonitrile (ACN) and 0.1% Formic acid (FA). Buffer B was 95% ACN and 0.1% FA.

For comparison between biotin-NHS and biotin-SS-NHS samples, a 35-cm homemade analytical column with pre-column was used. Liquid chromatography gradient was at a flow of 300 nL/min using the following 67-minute gradient profile: (min:%B) 0:3, 3:8, 40:27, 52:42, 53:90, 60:90, 67:100. Top 12 method with a full-scan MS spectrum with mass range of 350-1660 m/z was collected at a resolution of 120000, maximum injection time of 30 ms, and an AGC target of 2e5, MS/MS scan was acquired at 15000 resolution, maximum injection time of 60 ms, and an AGC target of 2e5. Normalized collision energy (NCE) was set to 28. Dynamic exclusion was set to 30 s. Charge state exclusion was set to ignore unassigned, +1, +5 and greater charges. Comparison between biotin-NHS and unlabeled samples, and comparison between inclusion and exclusion of singly-charged peptides were done on a 50-cm µPAC column, with and without pre-column, respectively. Liquid chromatography gradient was at a flow of 300 nL/min using the following 85-minute gradient profile: (min:%B) 0:4, 5:9, 10:10, 15:12, 20:14, 25:15, 30:17, 35:18, 40:19, 45:21, 50:24, 55:27, 60:80, 85:80. Top 12 method with a full-scan MS spectrum with mass range of 400-1800 m/z was collected at a resolution of 60000, maximum injection time of 75 ms, and an AGC target of 3e6. MS/MS scan was acquired at 15000 resolution, maximum injection time of 50 ms, and an AGC target of 5e4. NCE was set to 28. Dynamic exclusion was set to 20 s. Charge state exclusion was set to ignore unassigned, +1, +5 and greater than +8 charges, but +1 was removed from the list during the inclusion of singly-charged peptides.

### Labeling efficiency analysis

Fluorescence signal was measured from biotinylated peptides (biotin-NHS or biotin-SS-NHS) using Quantitative Fluorometric Peptide Assay (Thermo Scientific Pierce, cat.no.: 23290) and following the manufacturer’s protocol. An unlabeled peptide sample and 0.1% FA in water (blank) were used as controls for minimum and maximum labeling, respectively. To confirm peptide amounts from the fluorometric assay were comparable, peptide totals were measured using Quantitative Colorimetric Peptide Assay (Thermo Scientific Pierce, cat. no.: 23275).

### Data processing and analysis

Raw MS DDA data acquired from the Q Exactive HF were searched with MaxQuant (version 1.6.2.10) using the built-in Andromeda search engine, and embedded standard Orbitrap settings which included first search peptide tolerance at 20 ppm and main search peptide tolerance at 4.5 ppm. The false discovery rate for protein, peptide, and PSM were set at 1%. Trypsin/P specific digestion mode was used. Carbamidomethyl (C) was set as fixed modification, but was set as variable modification during the assessment of carbamidomethyl cysteines.. Oxidation (M) and Acetyl (Protein N-term) were set as dynamic modification. Additional dynamic modifications, as described in Supplementary Figure 1, were assigned for lysine and N terminus, as needed. For the comparison between biotin-NHS and biotin-NHS-SS, raw files for each biotinylated samples (n=5 for biotin-NHS and biotin-SS-NHS) were searched separately but grouped with unlabeled controls (n=5). The human protein database was downloaded from Uniprot (2018_01; 20,245 sequences). For the comparison between biotin-NHS (n=5) and unlabeled controls (n=5), and comparison between inclusion (n=5) and exclusion (n=5) of singly-charged peptides, the human protein database was downloaded from Uniprot (2020_03; 20,365 sequences). Common contaminants were embedded from MaxQuant. MaxQuant peptide identifications were obtained from the evidence.txt file output. Peptide sequences that were labeled as “Potential contaminant” or “Reverse” were excluded from further analysis. Determination of labeling efficiency and other comparison of features were done using R. All statistical tests were done using t-tests. For the statistical testing of the amino acid frequencies at the N terminus, an adjusted p-value was calculated using Benjamini and Hochberg method. All box plots prepared were in the style of Tukey: the box boundaries represent Q1 and Q3 while the line within the box represents the median; whiskers extend from the upper and lower quartiles to the maximum and minimum values, respectively, with the outliers excluded. Outliers are defined as 1.5 × IQR of Q1 or Q3 and are displayed as points beyond the minimum and maximum values, respectively.

## RESULTS

### Comparison biotin-NHS and biotin-SS-NHS

Tryptic peptides were prepared from HeLa lysates and were labeled with either biotin-NHS or biotin-SS-NHS. During biotinylation, primary amines (—NH_2_) at the N terminus or lysine residues makes a nucleophilic attack on the NHS ester moiety present in the biotin labels. This yields in a stable amide bond linking biotin and the peptide. To confirm biotinylation, we assessed the relative number of free amino groups in the peptides by conducting an assay using an amine-reactive fluorescent agent. In this assay, biotinylated amino groups are blocked from reacting with the fluorescent agent. Therefore, a fully biotinylated sample would yield a lower fluorescence signal. We found that both biotinylating reagents effectively labeled amine groups in the samples, with labeling efficiencies observed to be greater than 90% (Figure 1A).

**Figure 1.**
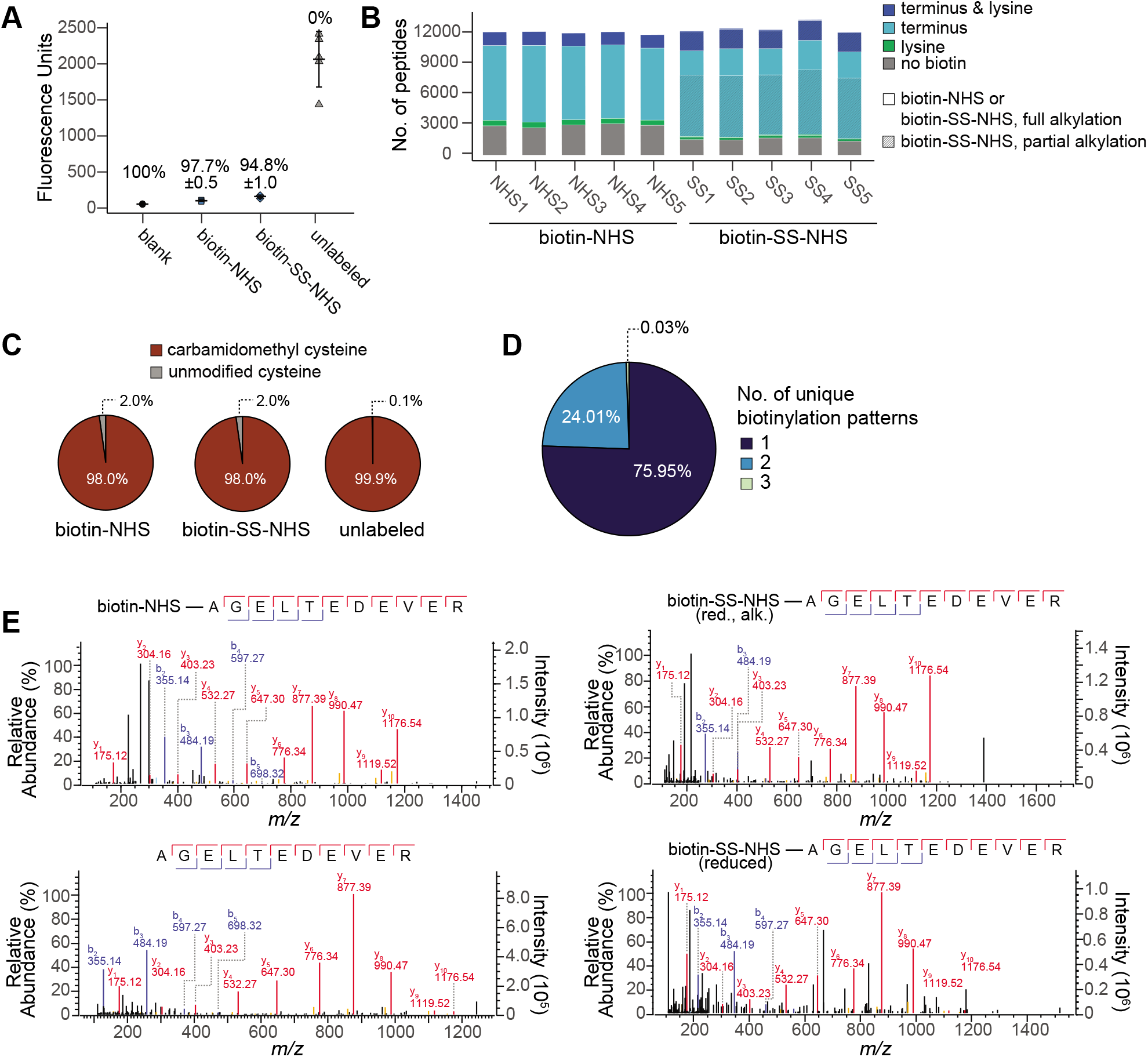
Comparison between biotin-NHS and biotin-SS-NHS peptides. **(A)** Biotin labeling efficiency using a fluorometric peptide assay (n=5 technical replicates for blank, biotin-NHS, and unlabeled; n=4 technical replicates for biotin-SS-NHS). Mean fluorescence units ± standard deviation plotted as points. Calculated labeling efficiency is displayed for each sample as mean ± standard deviation. **(B)** Comparison of peptide identifications between biotin-NHS and biotin-SS-NHS samples during LC-MS/MS analysis (n=5 technical replicates). Stacked bar plots represent total number of peptides detected in each sample broken down by the distribution of biotin labels between reaction sites within a peptide. Bar plots for the biotin-SS-NHS samples are further broken down based on the reduction and alkylation state. **(C)** Carbamidomethyl modification of cysteine residues. Each pie chart represents the average proportion of carbamidomethyl cysteine in the samples (n=5 technical replicates). **(D)** Biotinylation patterns in biotin-SS-NHS samples. The pie chart shows the average proportion of fully biotinylated peptides that have 1, 2, or 3 unique biotinylation patterns in the biotin-SS-NHS sample. **(E)** Representative mass spectra of the same peptide in biotin-NHS, biotin-SS-NHS and unlabeled samples. For the biotin-SS-NHS sample, the mass spectra for the peptide with a reduced and alkylated label is shown.

We then examined peptide biotinylation by LC-MS/MS. We observed that there was a similar number of total peptide identifications between the two biotin labels (Figure 1B). However, in general, we found slightly more biotinylated peptides in the biotin-SS-NHS samples compared to the biotin-NHS samples. On average, 88% of peptides were biotinylated in biotin-SS-NHS samples, whereas 76% of peptides were biotinylated in biotin-NHS samples. In terms of amine groups—i.e., available labeling sites, we observed an even split between unlabeled and biotinylated amino groups after LC-MS/MS analysis. On the other hand, 63% of amines in biotin-SS-NHS samples were biotinylated (Supplementary Figure 1).

Unlike biotin-NHS, biotin-SS-NHS occurred in non-reduced (<1% biotinylated termini and lysins), reduced (71% biotinylated termini; 87% biotinylated lysins) and alkylated (29% biotinylated termini; 13% biotinylated lysins) forms (Figure 1B, Supplementary Figures 2, 3). To test if the observed incomplete alkylation is specific to the biotin or if the alkylation conditions were not optimal, we determined the reduction and alkylation efficiency of cysteine residues. Here we observed that carbamidomethyl modification of cysteine residues was ≥98% in all samples (Figure 1C). We further found that 24% of fully biotinylated peptides in biotin-SS-NHS samples occurred in multiple forms (Figures 1D, E). This suggests that the biotin-SS-NHS label is not as readily alkylated under the same conditions as cysteine residues. The occurrence of peptides in multiple modification states was particularly worrying as it would likely reduce quantitative precision. High and consistent alkylation levels could possibly be achieved by optimizing conditions, but this was beyond the scope of this manuscript.

### Characteristics of biotinylated peptides

Since biotin-SS-NHS appeared to be incompletely and inconsistently alkylated, we decided to continue our studies using the biotin-NHS label to reduce search complexity and to have a more precise quantification of biotinylated peptides.

We next determined if the detection of biotin-NHS labeled peptides was comparable to normal unlabelled tryptic peptides. Using identical LC gradient and MS settings we found that fewer biotin-NHS than unlabeled peptides were detected (Figure 2A, Supplementary Figure 4). Unlike for biotin-SS-NHS this discrepancy could not be attributed to the fragment ion coverage (Figure 2B). We found that the b and y fragment ion coverage from both unlabeled and biotinylated samples were comparable. We also noted that the database (Andromeda) scores for the peptides detected from both groups were comparable, with median scores at 85.4 and 76.2 for biotinylated and unlabeled samples, respectively (Supplementary Figure 5). This suggests that our observations for the fragment ion coverage is not a result of having poorer signal-to-noise ratio in biotinylated samples. In addition, we observed that while there were fewer biotinylated peptides detected, the intensities of these peptides were comparable to unlabeled peptides (Figure 2C).

**Figure 2.**
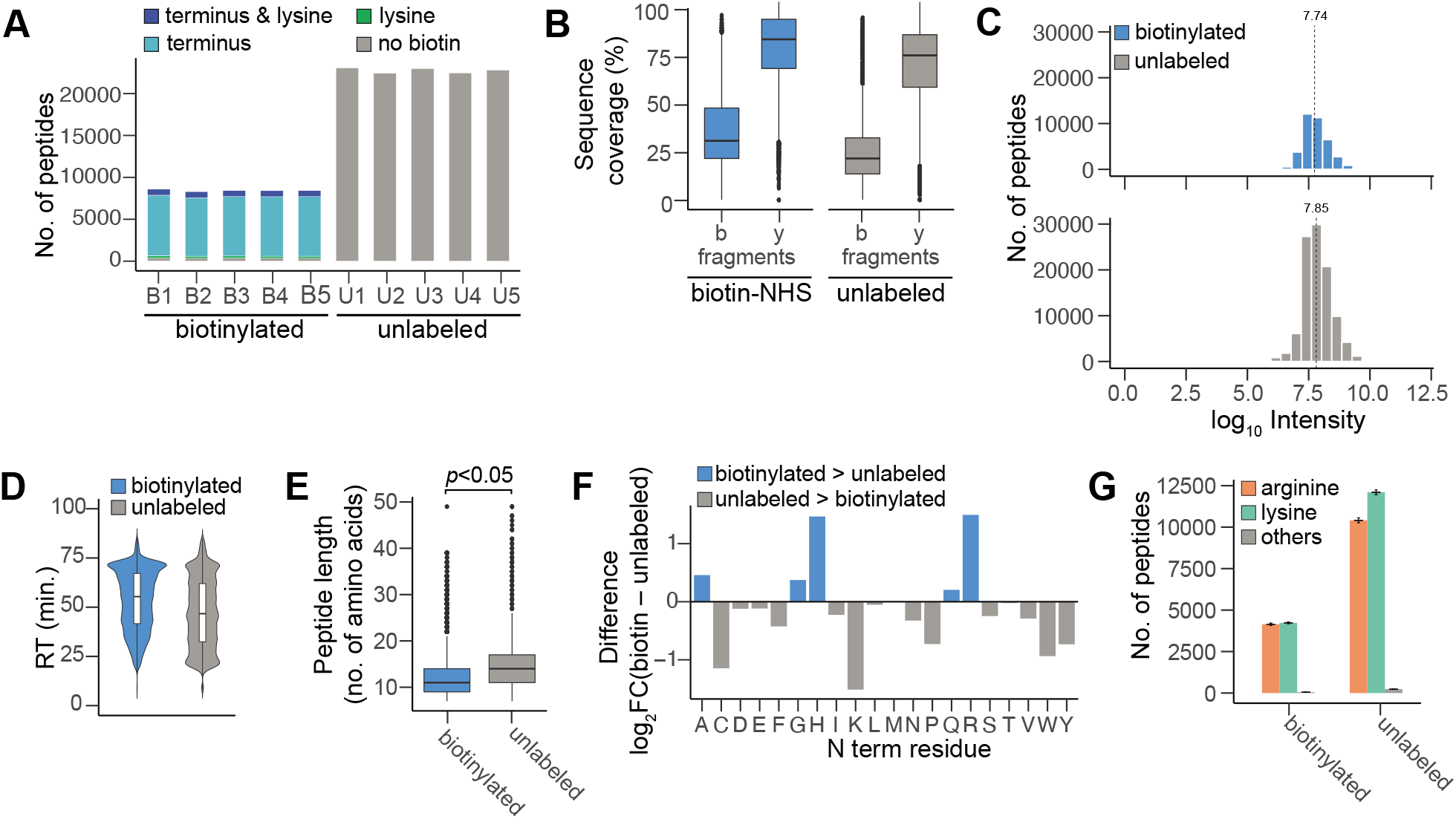
Comparison between biotin-NHS peptide and unlabeled peptides. **(A)** Comparison of peptide identifications between biotin-NHS samples and unlabeled samples (n=5 technical replicates). Stacked bar plots represent total number of peptides detected in each sample broken down by the distribution of biotin labels between reaction sites within a peptide. **(B)** Sequence coverage of b and y fragment ions in biotin-NHS (blue) and unlabeled (gray) samples (n=5 technical replicates). Tukey style box plots show median sequence coverage. **(C)** Distribution of peptide intensity in the biotinylated samples (top) and unlabeled samples (bottom). The dashed line indicates the median log_10_ Intensity. **(D)** Comparison of the observed retention time in biotinylated peptides and unlabeled peptides (n=5 technical replicates). Tukey style box plot is superimposed over a violin plot. The box plot shows the median retention time. **(E)** Comparison of peptide lengths observed in the biotinylated (blue) peptides and unlabeled (gray) peptides (n=5 technical replicates). Tukey style box plots show median peptide length. **(F)** Comparison of amino acid frequency in the N terminal position of biotinylated and unlabeled peptides (n=5 technical replicates). Bar plots represent the average difference between the log_2_FC calculated for each amino acid in each sample. FC is the ratio between the observed frequency of the amino acid at the N terminus and the expected amino acid frequency in the human proteome. Bar plots were annotated based on which group the amino acid has a higher log_2_FC (blue for biotinylated; gray for unlabeled). **(G)** Comparison of amino acid frequency in the C terminal position of biotinylated and unlabeled peptides (n=5 technical replicates). Bar plots indicate the average ± standard deviation of total number of peptides that either have arginine (orange) or lysine (green) at the C terminus in each group. Peptides that do not terminate with either arginine or lysine are in “others” (gray).

To elucidate reasons for the low biotinylated peptide detection, we characterized and compared the peptides detected in the unlabeled samples against those in the biotin-NHS samples. We initially inspected the retention time (RT) of the peptides and found that biotinylated peptides were likely to elute later (Figure 2D). Furthermore, this delay in RT was exacerbated as additional labels were attached to the peptide (Supplementary Figure 6). We also found that biotinylated peptides were significantly shorter (*p*<0.05) compared to unlabeled peptides. On average, biotinylated peptides were 12 amino acids long whereas unlabeled peptides were 14 amino acids (Figure 2E).

Examination of the N terminal residue revealed that the biotinylated peptides had different amino acid frequency compared to unlabeled peptides. We found that the biotinylated samples had a significant increase (*adj. p*<0.05) in the proportion of arginine (mean difference=1.5), histidine (mean difference=1.5), glycine (mean difference=0.4), glutamine (mean difference=0.2), and alanine (mean difference=0.5) at the N terminus (Figure 2F). For all other amino acids, except methionine and threonine, we found a significant decrease instead.

Stereochemistry potentially explains the over-representation of glycine and alanine in the biotinylated samples. Glycine has the smallest side chains out of all the amino acids, while alanine has the smallest hydrophobic side chain. The increase in the proportion of glycine and alanine in the biotinylated samples, suggests that these amino acids were often observed at the N terminus of biotinylated peptides because the side chains might render the primary amine more accessible.

We hypothesize that the overrepresentation of histidine and arginine in the biotinylated samples relates to the contribution of these amino acids’ side chains to the overall charge of the peptides. The reaction of primary amines in the N terminus and lysine side chains with the biotin label reduced the overall positive charge in the peptide. As a possible consequence, peptide detection becomes less permissible as typical mass spectrometric settings exclude uncharged or singly-charged entities for further analysis. Our observation suggests that there may be a positive selection towards biotinylated peptides with a positive selection towards biotinylated peptides with a positively-charged histidine or arginine.

We also examined the C terminal residue. Due to the use of trypsin during sample preparation, peptides that were detected typically had a lysine or arginine as its C terminal residue. We found that 53% of the unlabeled peptides had a C terminal lysine whereas 46% had a C terminal arginine. This distribution was altered in the biotinylated samples where peptides that have a C terminal lysine were reduced to 50% and peptides with a C terminal arginine increased to 49% (Figure 2G). Furthermore, we observed that peptides that were fully biotinylated disproportionately had arginine as its C terminal residue (Supplementary Figure 7). In fully biotinylated peptides, only 10% peptides had a lysine in the C terminus. The shift in the proportion of C terminal residues further supports the hypothesis that peptides with biotinylation are losing positive charges which made detection less permissible. This is not to suggest that fully biotinylated peptides that terminate with a lysine are completely undetectable; these peptides were observed in our data. However, these peptides were rare. In addition, these peptides possessed characteristics that were typical of biotinylated peptides, as described above. Most notable of which is the over-representation of histidine as a compensatory mechanism for the charge losses incurred during labeling.

### Optimization of biotinylated peptide detection

Since the biotinylation of peptide demonstrably affected its detection during LC-MS/MS analysis, we sought to determine if additional interventions can improve it.

For normal tryptic peptides one can expect most peptide precursors to carry at least two charges and singly charged precursors will predominantly originate from non-peptide contaminants. We hypothesized that the elimination of charges by biotinylation will result in more singly charged peptide precursors. We further hypothesized that the increase in detected biotinylated peptides following inclusion of singly charged precursors in the selection for fragmentation will outweigh the loss associated with fragmentation of non-peptide contaminants.

We found that the number of singly-charged peptides increased in both biotinylated and unlabeled samples after altering the MS settings. However, we found that the biotinylated samples had significantly more singly-charged peptides compared to unlabeled samples in both settings (Supplementary Figures 8A, 8B). In all, we indeed found a modest average 21% increase in peptide identifications when we permitted the analysis of singly-charged precursors (Figure 3A, Figure 3B). In contrast, the smaller net gain in identifications we observed in the unlabeled sample only translated to a 7% increase (Supplementary Figure 8C). Furthermore, we observed that the retention time of the singly-charged peptides were more evenly distributed throughout the gradient, whereas higher charged precursors where skewed towards the hydrophobic end of gradient (Figure 3B). Meaning, this alteration in the MS settings is possibly further enhanced by the liquid chromatography in that singly-charged peptides better separated on standard gradients and made more amenable for detection, especially in a DDA setting. As expected we also observed a modest decline of on average 12 and 18% in doubly- and triply-charged peptides detected, respectively (Figure 3C). Overall, we observed a net gain of 21% in biotinylated peptide identifications showing that the gain in singly-charged peptides outweighs the losses from higher charge states. In-depth analysis of the singly-charged peptides revealed that the majority of these are biotinylated at the N terminus (Figure 3D). The increase in N terminally labeled peptides may be of particular value in certain studies that employ biotin handles to enrich select protein amino termini to examining proteolytic activity.

**Figure 3.**
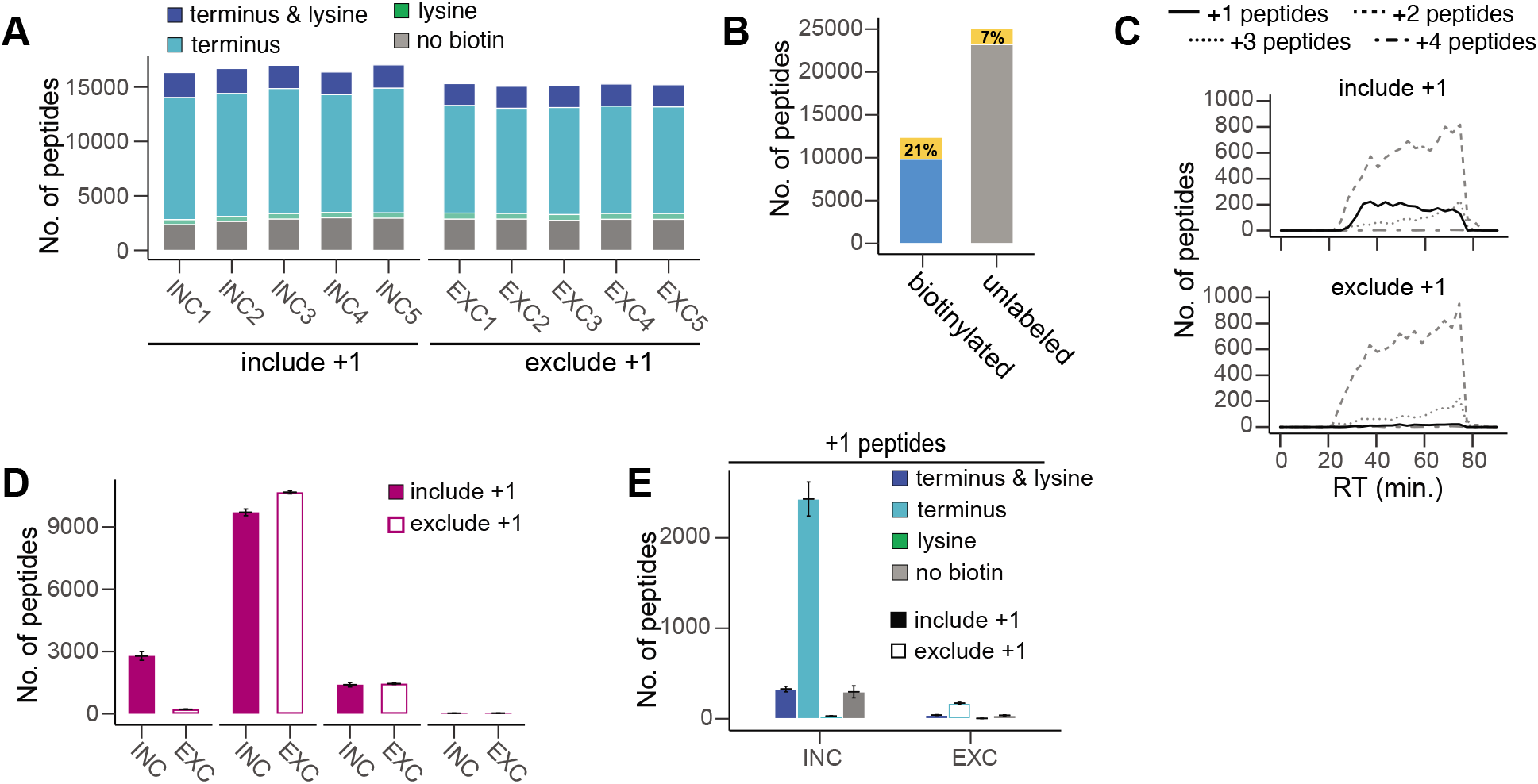
Optimization of biotinylated peptide detection. **(A)** Comparison of peptide Identification with inclusion and exclusion of singly-charged peptides (n=5 technical replicates). Bar plots represent total number of peptides detected in each sample. Additional annotation is available to provide details on the distribution of biotin labels between potential locations within a peptide. **(B)** Comparison of the representative elution profiles observed with either inclusion and exclusion of singly-charged peptides. The line plots represent the number of peptides eluting at a particular retention time. Peptides are stratified based on charge. **(C)** Charge distribution of the peptides detected with inclusion or exclusion of singly-charged peptides (n=5 technical replicates for exclude singly-charged peptides, n=2 for include singly-charged peptides). Bar plots show average number ± standard deviation of peptides detected within each type of peptide charge. **(D)** Biotinylation of singly-charged peptides with inclusion or exclusion singly-charged peptides. Bar plots represent the average number ± standard deviation of singly-charged peptides.

## DISCUSSION

Although commercially-available biotin labels have wide utility in proteomics, our studies revealed that attachment of this label to peptides affected peptide detection by LC-MS/MS. The use of anti-biotin antibodies^7,10^ can support the use of non-cleavable biotin that showed more favourable peptide separation, ionization and fragmentation behaviour. Yet, overall, biotinylated samples yielded fewer peptide identifications compared to unlabeled samples. Recent work has nicely demonstrated that fragmentation of biotinylated peptides yields signature fragment ions that can be utilized for more reliable detection of partially biotinylated peptides^11^. This work is critically useful in the reliable identification of biotinylated peptides, especially in experiments wherein the spectra of biotinylated peptides are rare and buried in the spectra of unlabeled counterparts.

However, detection and use of signature peptides requires efficient separation, ionization and fragmentation of biotinylated peptides to occur. Our work provides possible explanations for low identification rates of, in particular highly-biotinylated peptides, and points out possible mitigation strategies. The difference observed in the biotinylated samples can be attributed to the quenching of the positive charge in the primary amines located in the N terminus and side chains of the lysine residues. The loss of this positive charge applied selective favour towards detection of peptides that were characteristically different compared to unlabeled tryptic peptides. We observed that biotinylated peptides were shorter and had later retention times, despite their intensities and fragment ion coverage appearing to be comparable to unlabeled controls. Additionally, amino acids at both terminal regions were markedly different between biotinylated and unlabeled peptides. We found that the N terminus was more likely to be biotinylated if the terminal residue was an alanine or glycine. This is possibly because these amino acids present the least amount of steric hindrance. We also noted that the N terminus and C terminus of biotinylated peptides had amino acids that are able to compensate for the charge losses resulting from labeling. In the C terminal region, we found biotinylated peptides—especially those that are completely labeled— to disproportionately have arginine, as opposed to lysine. In the N terminal region, both histidine and arginine were found to be overrepresented.

The described effects of biotinylation on the detection of peptides is crucial to address in proteomic workflows, especially because the biotinylated peptide provides the most direct evidence for the target protein and site of biotinylation. Here, we demonstrated that inclusion of singly-charged peptides could provide modest increase in the number of peptide identifications. We further demonstrated a shift in the retention time of biotinylated peptides suggesting that the liquid chromatography gradient needs to be optimized. Specifically, allowing for a shallower gradient at higher organic solvent concentrations may be necessary in order to fully separate biotinylated peptides. We conclude that biotinylation strongly affects the chromatographic and ionization/fragmentation properties of peptides. Optimized method parameters may compensate some of the negative effects but in particular high or fully modified peptides remain challenging to detect.

## ACKNOWLEDGMENTS

This work was partially supported by grants from the Michael Cuccione Foundation, the BC Children’s Hospital Foundation (to P.F.L.) and Canadian Institutes of Health Research (PJT-169190 to P.F.L.). L.N. was supported by a fellowship from the Michael Cuccione Childhood Cancer Research Program. P.F.L. was supported by the Canada Research Chairs program and the Michael Smith Foundation for Health Research Scholar program.

---

**Figure S1.**
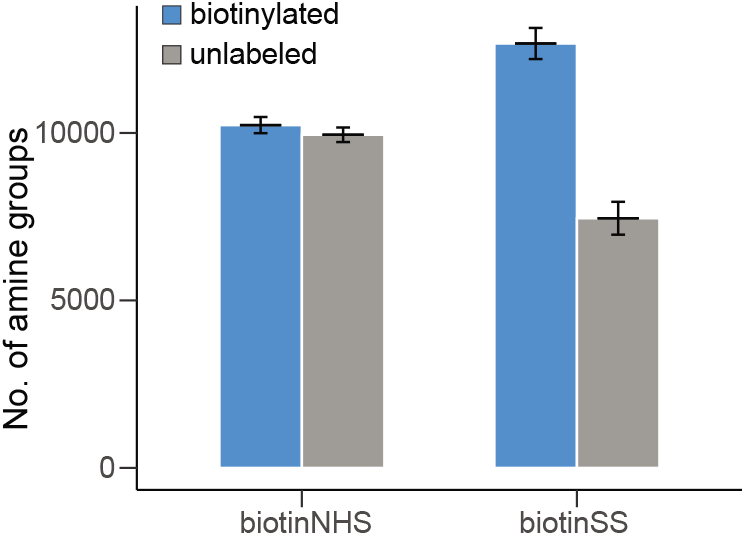
Labeling of amine groups observed in the biotinylated samples after LC-MS/MS analysis (n=5 technical replicates). Bar plots represent the average number ± standard deviation of amine groups that were biotinylated (blue) or unlabeled (gray) within each group.

**Figure S2.**
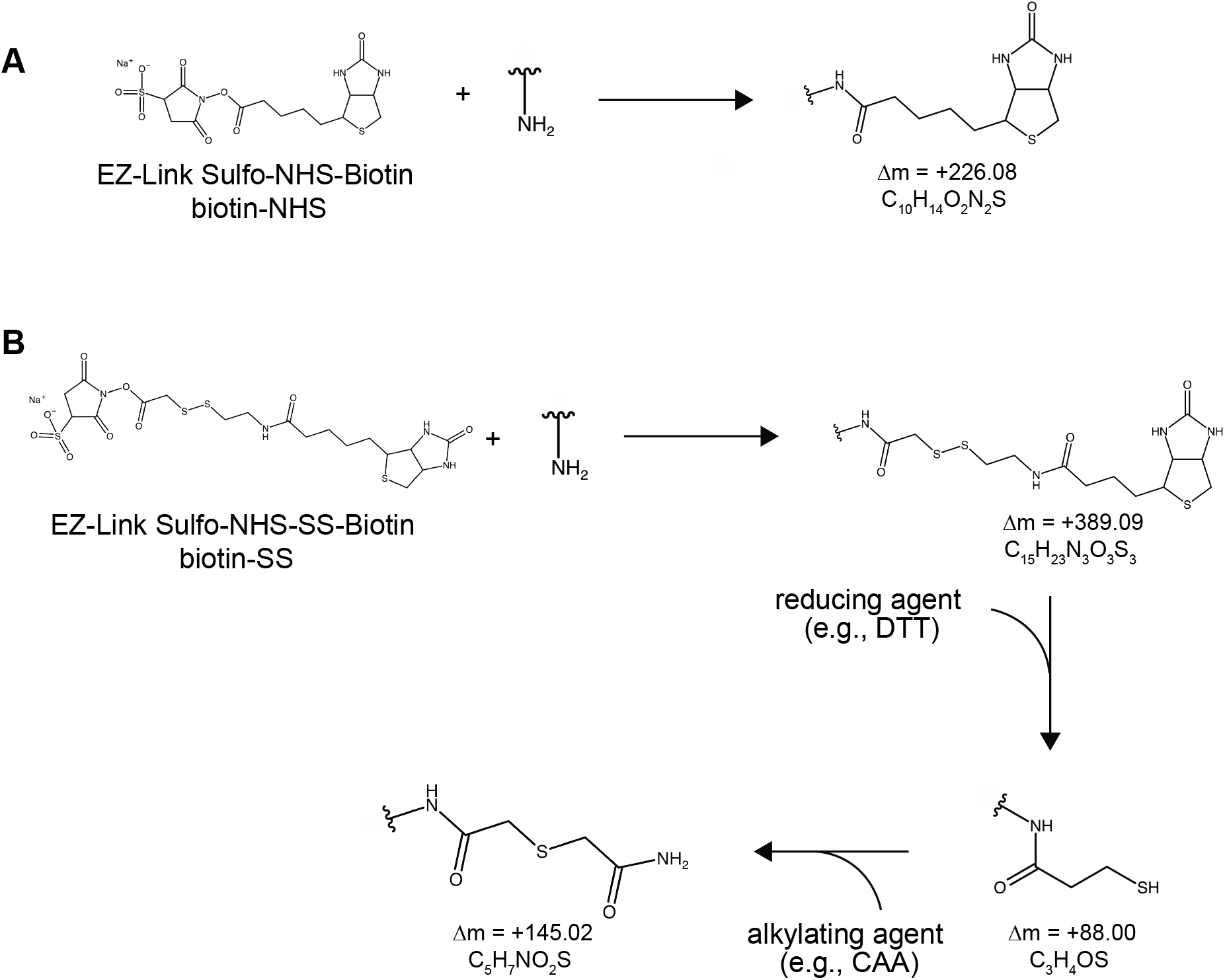
Summary of expected modifications from biotin labeling. **(A)** The reaction between EZ-Link Sulfo-NHS-Biotin (biotin-NHS) with a primary amine and the expected modification is shown. **(B)** The reaction between EZ-Link Sulfo-NHS-SS-Biotin (biotin-SS-NHS) with a primary amine and the expected modification is shown. Other possible modifications resulting from reduction and alkylation of the biotin label are also shown.

**Figure S3.**
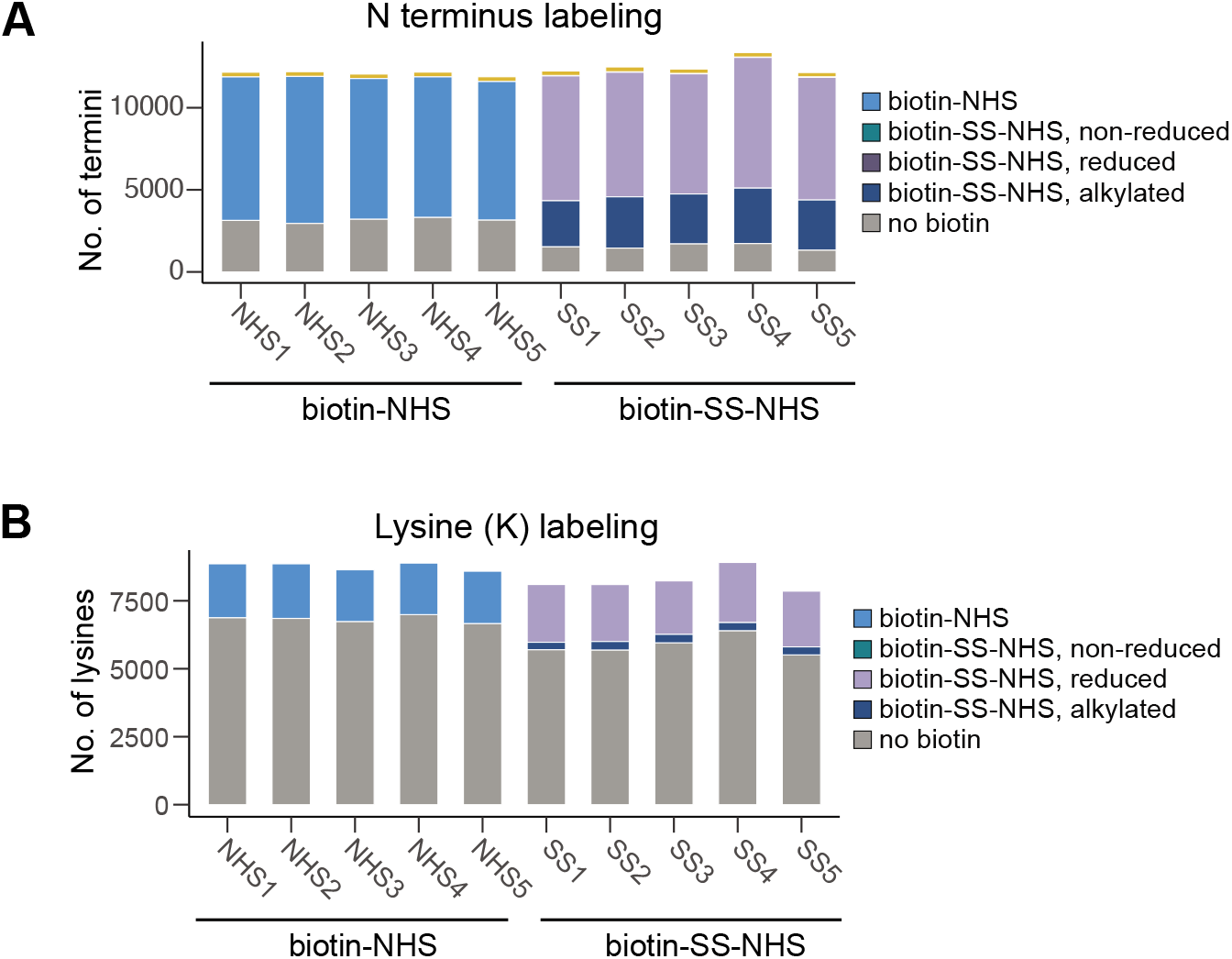
N-term and lysine biotinylation (n=5 technical replicates). **(A)** Bar plots represent the number of termini available in each sample. Non-gray portions of the bar represent the proportion of biotinylated termini in each sample. For biotin-SS-NHS samples, an additional level of annotation is added by indicating which form of the label is found. **(B)** Bar plots represent the number of lysines available in each sample. Non-gray portion of the bar represents the proportion of biotinylated lysine in each sample. For biotin-SS-NHS samples, an additional level of annotation is added by indicating which form of the label is found.

**Figure S4.**
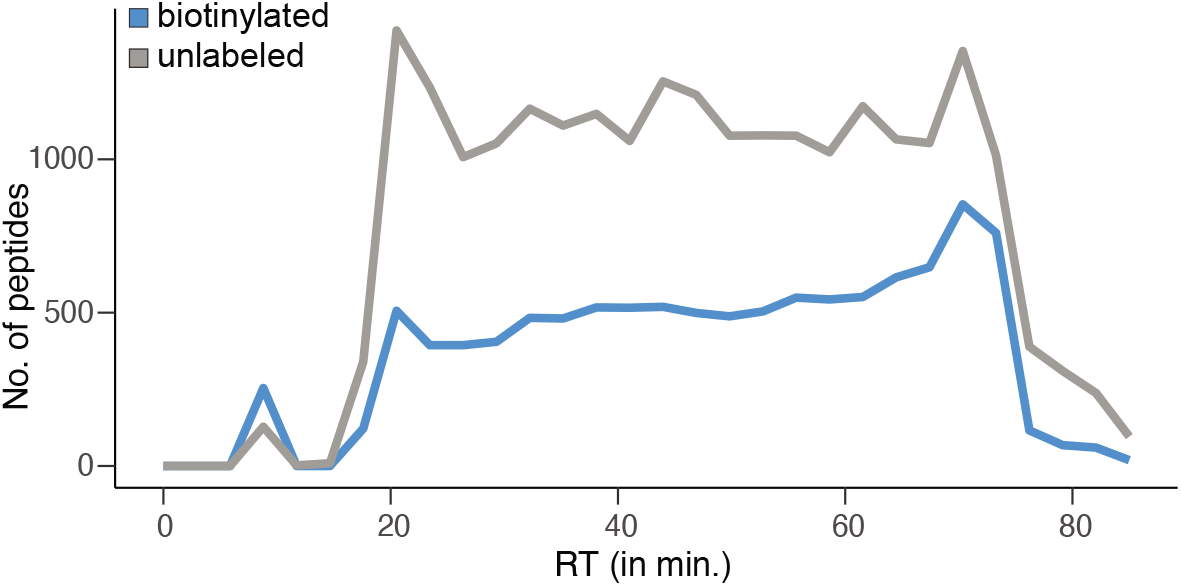
Representative elution profile of unlabeled and biotinylated peptides in the same liquid chromatography gradient. The line plots indicate the number of peptides eluting at a particular retention time.

**Figure S5.**
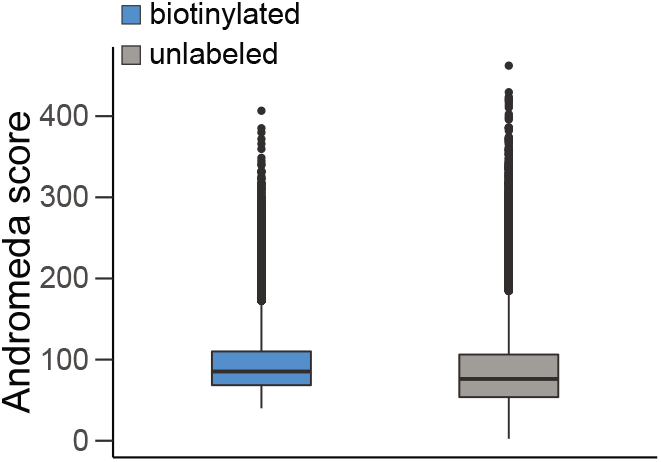
Database scores for the peptides identified in the biotinylated and unlabeled samples (n=5 technical replicates). Tukey style boxplot show distribution of peptide Andromeda scores within each experimental group.

**Figure S6.**
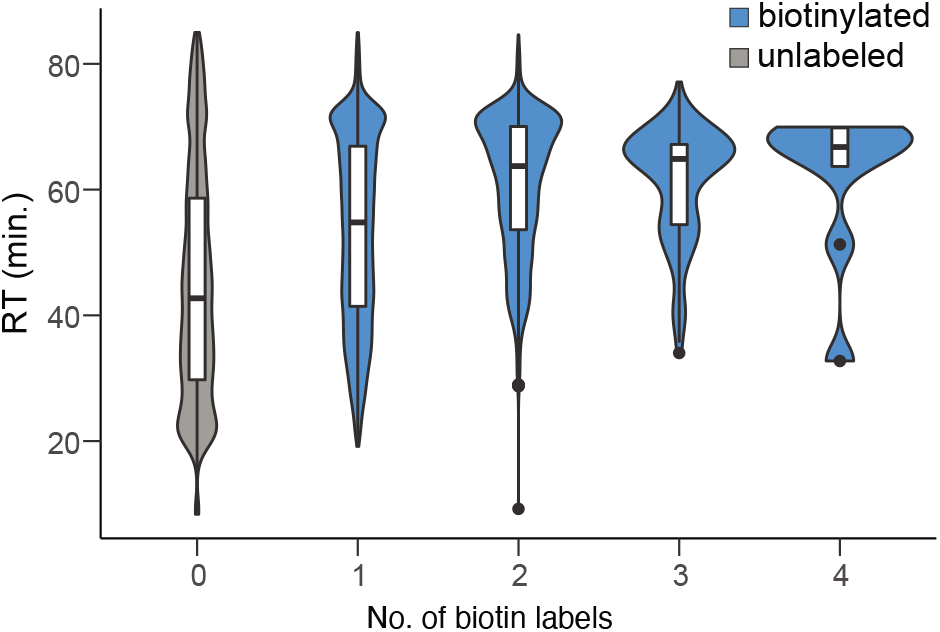
Retention time of peptides based on the number of biotin labels attached (n=5 technical replicates). Tukey style box plots are superimposed over violin plots. Violin plot indicates density (wider areas indicate more peptides) at a particular retention time. The box plots show median retention time within each group.

**Figure S7.**
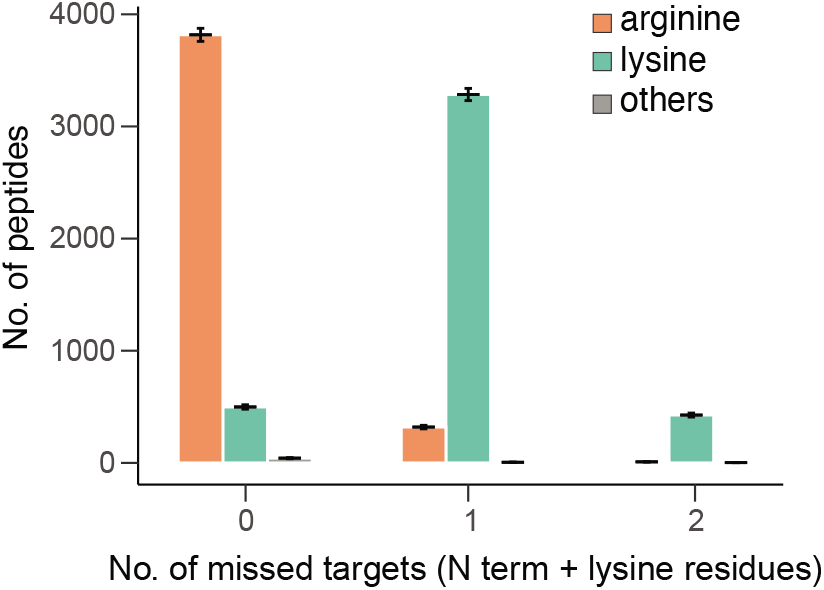
C terminal residue based on number of missed targets. Bar plots represent the average number ± standard deviation of peptides that either have an arginine (orange) or lysine (green) at the C terminus. Other C terminus amino acids were all grouped under “others” (gray). Number of missed targets refer to the number of N terminus and/or lysine residues that remained unlabeled within a particular peptide.

**Figure S8.**
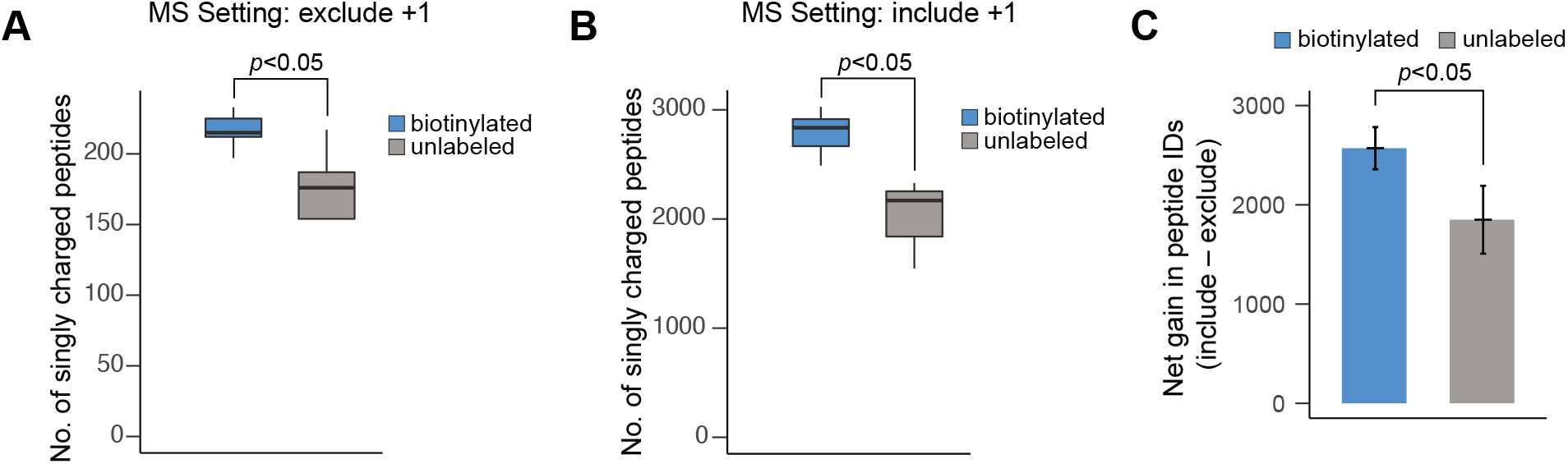
Singly-charged peptides identified in biotinylated and unlabeled samples (n=5 technical replicates). **(A)** Tukey style box plots show median number of singly-charged peptides identified in the biotinylated (blue) and unlabeled (gray) samples when the MS setting excludes singly-charged precursors. **(B)** Tukey style box plots show median number of singly-charged peptides identified in the biotinylated (blue) and unlabeled (gray) samples when the MS setting includes singly-charged precursors. **(C)** Net increase in singly-charged peptide identifications from both experimental groups. Bar plots represent the average ± standard deviation increase in the number of peptide identifications in biotinylated (blue) and unlabeled (gray) samples.

## Notes

### Competing Interest Statement

The authors have declared no competing interest.

